# Inactivation of SARS-CoV-2 on surfaces and in solution with Virusend (TX-10), a novel disinfectant

**DOI:** 10.1101/2020.11.25.394288

**Authors:** Enyia R. Anderson, Grant L. Hughes, Edward I. Patterson

**Affiliations:** Departments of Vector Biology and Tropical Disease Biology, Centre for Neglected Tropical Disease, Liverpool School of Tropical Medicine, Liverpool L3 5QA, UK

## Abstract

Until an effective vaccine against SARS-CoV-2 is available on a widespread scale, the control of the COVID-19 pandemic is reliant upon effective pandemic control measures. The ability of SARS-CoV-2 to remain viable on surfaces and in aerosols, means indirect contact transmission can occur and so there is an opportunity to reduce transmission using effective disinfectants in public and communal spaces. Virusend (TX-10), a novel disinfectant, has been developed as a highly effective disinfectant against a range of microbial agents. Here we investigate the ability of Virusend (TX-10) to inactivation SARS-CoV-2. Using surface and solution inactivation assays, we show that Virusend (TX-10) is able to reduce SARS-CoV-2 viral titre by 4log_10_ PFU/mL within 1 minute of contact. Ensuring disinfectants are highly effective against SARS-CoV-2 is important in eliminating environmental sources of the virus to control the COVID-19 pandemic.

## Introduction

Severe acute respiratory syndrome coronavirus 2 (SARS-CoV-2) is a novel coronavirus that is the causative agent of COVID-19 which first emerged in late 2019 [1]. Countries are working to control transmission of SARS-CoV-2 with the ultimate goal of production and large-scale manufacture of an effective vaccine [2-4]. However, until an effective vaccine is found, control of the virus is limited to implementing measures such as contact tracing, quarantine, enforcing strict social distancing, advising frequent hand hygiene and infection control measures in hospital environments [5]. During the 2002 outbreak of SARS-CoV-1, and the 2012 Middle East respiratory syndrome-related (MERS)-CoV outbreak, virus stability facilitated transmission events [6]. Similarly, research has shown that SARS-CoV-2 can remain viable on surfaces, notably plastic and stainless steel for up to 72 hours post inoculation, and in aerosols for at least 3 hours, meaning effective disinfectants can prevent indirect contact transmission [7]. Virusend (TX-10) has been developed to work as a highly effective disinfectant that rapidly inactivates infectious enveloped viruses. As communities begin to reopen and people return to the workplace, effective and quick disinfection of communal areas is paramount to maintaining control of COVID-19. Here we present the evidence that Virusend TX-10 can reduce SAR-CoV-2 virus within one minute both in solution and on surfaces.

## Methods and Materials

### Cell culture and viruses

Vero E6 cells (C1008: African green monkey kidney cells), obtained from Public Health England, were maintained in Dulbecco’s minimal essential medium (DMEM) containing 10% foetal bovine serum (FBS) and 0.05 mg/ml gentamicin. Cells were kept at 37°C with 5% CO_2_. Passage 4 or 5 of SARS-CoV-2 isolate (REMRQ0001/Human/2020/Liverpool) from a clinical sample was used to assess inactivation of TX-10. On the fourth and fifth passages the virus was cultured in Vero E6 cells maintained in DMEM with 4% FBS and 0.05mg/mL gentamicin at 37°C and 5% CO_2_ as previously described [8]. The fifth passage of the virus was harvested 48 hours after inoculation and concentrated by passage through a centrifugal column (Amicon Ultra-15 100kDa MWCO). Virus was used immediately after concentrating.

### Virus Inactivation

Inactivation on surfaces were preformed using either 9.8log_10_ or 7.9log_10_ PFU/mL of SARS-CoV-2. Surface inactivation was carried out by inoculating stainless-steel discs with 50µL of virus and allowed to air dry at room temperature for 1 hour. Dried inoculum was incubated with 100µl of Virusend (TX-10; Pritchard Spray Technologies, Colchester, UK) or autoclaved water for either 30 seconds or 9.5 minutes, after which 900µL of DMEM containing 2% FBS and 0.05 mg/mL gentamicin was added and mixed until dried inoculum was dissolved. The sample was then transferred into a dilution series for virus quantification at exactly 1 minute or 10 minutes after addition of TX-10 to the dried inoculum. Solution inactivation assays used either 8.4log_10_ or 7.9log_10_ PFU/mL and were carried out by incubating 25μL of inoculum with 100μL of TX-10 or autoclaved water for either 1 minute or 10 minutes. After incubation 10mL of DMEM was added and transferred to a dilution series within 30 seconds of DMEM being added. All experiments were performed in duplicate.

### Cytotoxicity Assay

Cytotoxicity for surface inactivation was determined by inoculating stainless-steel discs with 50µL of DMEM containing 2% FBS and 0.05 mg/mL gentamicin and allowed to air dry at room temperature for 1 hour. Dried inoculum was incubated with 100µl of TX-10 or autoclaved water for 5 minutes, after which 900µL of DMEM containing 2% FBS and 0.05 mg/mL gentamicin was added and mixed until dried inoculum was dissolved. The sample was then transferred into a dilution series and a standard plaque assay performed. Cytotoxicity for solution assays were performed by incubating 25 µL of DMEM containing 2% FBS and 0.05 mg/mL gentamicin with 100 µL of TX-10 for 5 minutes, after which 10mL of DMEM was added and sample transferred to a dilution series for standard plaque assays. The cytotoxicity assays were performed in duplicate.

### Suppression Assay

Suppression for solution inactivation was assayed by adding 25μL of inoculum to 100μL of TX-10 in 10mL of DMEM and incubated for 30 seconds. After 30 seconds, the sample was transferred into a dilution series and a standard plaque assay preformed. The suppression assay was performed in duplicate.

### Virus Quantification and Viability

Samples from each condition were serial diluted 10-fold for quantification by standard plaque assay using Vero E6 cells. Cells were incubated for 72 hours at 37°C and 5% CO_2_, then fixed with 10% formalin and stained with 0.05% crystal violet solution. Plaques were counted to calculate virus titre. All samples were performed in technical duplicates.

## Results

For inactivation assays, Virusend TX-10 was directly placed on SARS-CoV-2 inoculum, for an incubation period of either 1 minute or 10 minutes. On the hard surface, contact time of 1 minute with Virusend TX-10 reduced SARS-CoV-2 titres to below the limit of detection for both high and low titre inoculum (Fig 1). A titre of 7.3log_10_ PFU/mL was recovered from the high titre, hard surface control samples. Similarly, incubation with Virusend TX-10 for 10 minutes reduced the virus titre to below the limit of detection, compared with 7.0log_10_ PFU/mL recovered from the high titre control. With a low titre inoculum, Virusend TX-10 also reduced SARS-CoV-2 titres to below the limit of detection after contact times of 1 and 10 minutes on hard surfaces. Titres of 5.3log_10_ PFU/mL and 5.9log_10_ PFU/mL were recovered from the 1- and 10-minute control samples, respectively. Cytotoxicity assays with Virusend TX-10 in the absence of virus were used to determine the limit of detection, the point at which Vero E6 cell death is due to the cytotoxicity of Virusend TX-10, and not virus. Cytopathic effect was observed to 3.0log_10_ PFU/mL (Fig. 1). Both inactivation and cytotoxicity assays confirm a reduction of at least 4.0log_10_ PFU/mL of infectious SARS-CoV-2 with high titre inoculum and a reduction of at least 2.3log_10_ PFU/mL with low titre inoculum (Fig 1).

**Fig 1.**
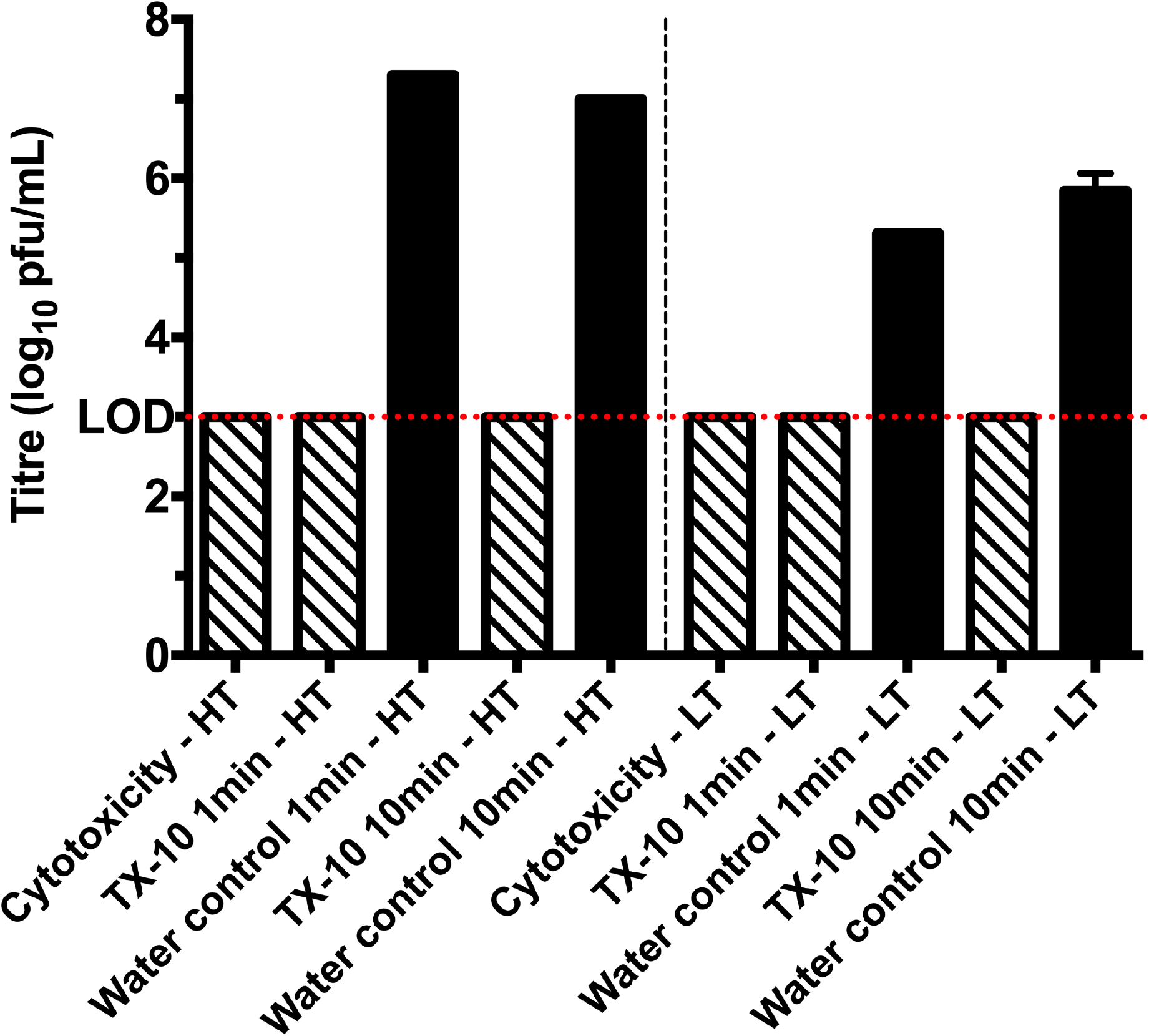
Virusend TX-10 reduces viral titre on hard surfaces by at least 4.0log_10_ PFU/mL with high titre (HT) viral inoculum after contact times of 1 minute and 10 minutes. When low titre (LT) inoculum was used, TX-10 reduces virus titre by at least a 2.31log_10_ PFU/mL at both 1 minute and 10-minute contact time. Diagonal pattern represents cytopathic effect caused by TX-10 and solid black represents the titre of infectious virus following each treatment. Limit of detection (LOD) (3.0log_10_ PFU/mL) is shown across the graph with a dotted red line.

For inactivation assays in solution, Virusend TX-10 was placed directly into solution with SARS-CoV-2 for either 1 or 10 minutes. An incubation period of 1 minute with Virusend TX-10 reduced the high titre inoculum from 6.00log_10_ PFU/mL, in the water control, to below the limit of detection (Fig 2A). A 10-minute incubation with Virusend TX-10 also reduced viral titre from 6.0log_10_ PFU/mL to below the limit of detection (Fig 2B). With the low titre inoculum, the addition of Virusend TX-10 reduced SARS-CoV-2 to below the limit of detection at both 1 minute and 10 minute incubation times (Fig 2). Titres of 5.6log_10_ PFU/mL were recovered from control samples at 1 minute and 10 minutes. A suppression assay for solution inactivation assays was used to demonstrate that dilution with 10mL of DMEM suppressed Virusend TX-10 inactivation of SARS-CoV-2 upon the completion of the assay. The addition of Virusend TX-10 to virus inoculum in 10mL of DMEM recovered a virus titre of 5.7log_10_ PFU/mL with high titre inoculum and 5.6log_10_ PFU/mL with low titre inoculum. Cytotoxicity assays for solution inactivation assays showed the limit of detection for these assays was 2.0log_10_ PFU/mL.

**Fig 2.**
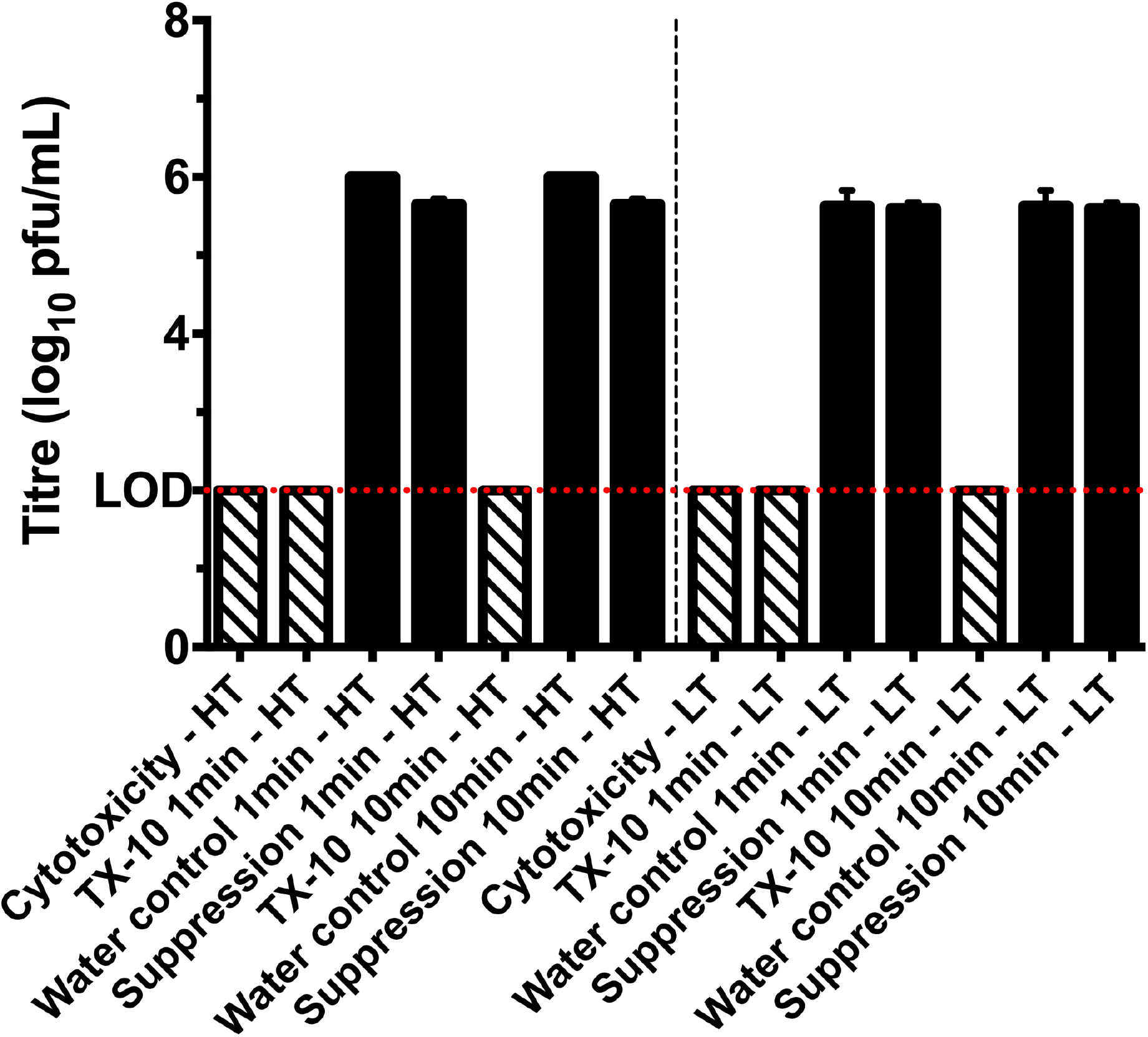
Virusend TX-10 reduces viral titre in solution by at least 4.0log_10_ PFU/mL when incubated with high titre (HT) virus inoculum for 1 minute and 10 minutes. When low titre (LT) inoculum was used, both incubation periods reduced the titre by at least 3.6log_10_ PFU/mL, to below the limit of detection. Diagonal pattern represents cytopathic effect caused by TX-10 and solid black represents the titre of infectious virus following each treatment. Limit of detection (LOD) (2.0log_10_ PFU/mL) is shown across the graph with a dotted red line.

## Discussion

SARS-CoV-2 can remain viable on surfaces, notably plastic and stainless steel for up to 72 hours post inoculation, and in aerosols for at least 3 hours [7]. In solutions, such as respiratory droplets, SARS-CoV-2 may remain viable for up to 14 days at 4°C, 7 days at room temperature, and for 1 to 2 days at 37°C [9]. Therefore, contaminated surfaces and solutions are a reservoir for transmission through fomites, meaning effective hygiene and environmental decontamination is crucial in helping to prevent the spread of COVID-19 [10, 11]. Disinfectant solutions of 75% ethanol and 10% sodium hypochlorite are able to reduce SARS-CoV-2 titre by at least 2.0log_10_ PFU/mL and 3.25log_10_ PFU/mL, respectively, within 5 minutes [9]. However, the WHO has recommended diluting household bleach 1:100 to reduce irritation to the user and contact times of 10 to 60 minutes to disinfect surfaces and when immersing items [12]. Rapid household disinfectants could reduce transmission in private residence and public spaces, such as offices. Here we have shown that Virusend TX-10 is able to reduce SARS-CoV-2 virus titre by at least 4.0log_10_ PFU/mL in 1 minute of contact time making it and effective disinfectant for households and public spaces.

An initial obstruction to the work presented here, was the need for a high virus titre to show a 4.0log_10_ PFU/mL reduction due to the cytotoxicity of Virusend TX-10 to Vero E6 cells. The limit of detection indicated the point at which cytopathic effect in Vero E6 cells is caused by Virusend TX-10 and not the virus. Therefore, to achieve a 4.0log_10_ PFU/mL reduction, SARS-CoV-2 had to be concentrated after harvesting to give stock titres of 8.4log_10_ and 9.8log_10_ PFU/mL. When a lower stock virus titre of 7.9log_10_ PFU/mL was used, a 4.0log_10_ PFU/mL reduction could not be demonstrated and would not meet the strict requirements of European Standard testing. However, these assays still showed a similar trend of inactivation. The use of higher viral titre in these assays indicates the effectiveness of Virusend TX-10, which may be necessary to inactivate SARS-CoV-2 in environments that are contaminated [13].

Disinfectants tested for use against other members of *Coronaviridae* have used surrogates, such as murine hepatitis virus, a lower biosafety level pathogen that can be grown to high titres and has structural and genetic similarities to SARS-CoV, to be able to carry out the assays more easily [14]. Surrogates are chosen to mimic the inactivation of the target virus, but the use of surrogates should be limited, and the target pathogen should be used when possible [15]. Here we have been able to test Virusend TX-10 against SARS-CoV-2, the causative agent of the current pandemic, therefore the ability of Virusend TX-10 to significantly reduce the viral titre of the relevant virus.

Virusend TX-10 is a detergent-based disinfectant. Other detergents, NP-40 and Triton X-100, have been shown to completely inactivate SARS-CoV-2 at a concentration of 0.5% [8]. Both Triton X-100 and NP-40 have also been shown to inactivated the small enveloped hepatitis C virus to below detectable levels, and Triton X-100 inactivates HIV-1 virus completely within 1 minute [16]. Like Virusend TX-10, the ability of detergents to inactivate harmful viruses mean they are important for disinfecting contaminated surfaces and solutions. The development of Virusend TX-10, and showing it is highly efficient at inactivating the circulating strain of SARS-CoV-2 specifically, is important to minimise community transmission of SARS-CoV-2.

Current advice focuses on increasing public engagement in essential control measures, such as high levels of hygiene in the home [17]. Virusend TX-10 can reduce the strain of demand on current hygiene product resources, to be used within private residences, communal public areas such as offices and hospital environments [18-20]. It can reduce viral titres on surfaces and in solution by at least 4.0log_10_ PFU/mL within 1 minute of contact making it highly suitable for rapid disinfection of private households and public spaces such as hospitals and offices. The development of disinfectants such as Virusend TX-10 and others is important as we continue efforts to reduce transmission of SARS-CoV-2.

## Acknowledgements

This work was supported by the Ministry of Defence. GLH was supported by the BBSRC (BB/T001240/1 and V011278/1), a Royal Society Wolfson Fellowship (RSWF\R1\180013), the NIH (R21AI138074), URKI (20197), and the NIHR (NIHR2000907). GLH is affiliated to the National Institute for Health Research Health Protection Research Unit (NIHR HPRU) in Emerging and Zoonotic Infections at University of Liverpool in partnership with Public Health England (PHE), in collaboration with Liverpool School of Tropical Medicine and the University of Oxford. GLH is based at LSTM. The views expressed are those of the author(s) and not necessarily those of the NHS, the NIHR, the Department of Health or Public Health England. EIP was supported by the Liverpool School of Tropical Medicine Director’s Catalyst Fund award.

## References

1. Wu F, Zhao S, Yu B, Chen YM, Wang W, Song ZG, et al. A new coronavirus associated with human respiratory disease in China. Nature. 2020;579(7798):265–9. Epub 2020/02/06. doi: 10.1038/s41586-020-2008-3. PubMed PMID: 32015508; PubMed Central PMCID: PMCPMC7094943.

2. Sharpe HR, Gilbride C, Allen E, Belij-Rammerstorfer S, Bissett C, Ewer K, et al. The early landscape of coronavirus disease 2019 vaccine development in the UK and rest of the world. Immunology. 2020;160(3):223–32. Epub 2020/05/28. doi: 10.1111/imm.13222. PubMed PMID: 32460358; PubMed Central PMCID: PMCPMC7283842.

3. Yamey G, Schaferhoff M, Hatchett R, Pate M, Zhao F, McDade KK. Ensuring global access to COVID-19 vaccines. Lancet. 2020;395(10234):1405–6. Epub 2020/04/04. doi: 10.1016/S0140-6736(20)30763-7. PubMed PMID: 32243778; PubMed Central PMCID: PMCPMC7271264.

4. Thanh Le T, Andreadakis Z, Kumar A, Gomez Roman R, Tollefsen S, Saville M, et al. The COVID-19 vaccine development landscape. Nat Rev Drug Discov. 2020;19(5):305–6. Epub 2020/04/11. doi: 10.1038/d41573-020-00073-5. PubMed PMID: 32273591.

5. Cheng VC, Wong SC, Chuang VW, So SY, Chen JH, Sridhar S, et al. The role of community-wide wearing of face mask for control of coronavirus disease 2019 (COVID-19) epidemic due to SARS-CoV-2. J Infect. 2020;81(1):107–14. Epub 2020/04/27. doi: 10.1016/j.jinf.2020.04.024. PubMed PMID: 32335167; PubMed Central PMCID: PMCPMC7177146.

6. Xiao S, Li Y, Wong TW, Hui DSC. Role of fomites in SARS transmission during the largest hospital outbreak in Hong Kong. PLoS One. 2017;12(7):e0181558. Epub 2017/07/21. doi: 10.1371/journal.pone.0181558. PubMed PMID: 28727803; PubMed Central PMCID: PMCPMC5519164.

7. van Doremalen N, Bushmaker T, Morris DH, Holbrook MG, Gamble A, Williamson BN, et al. Aerosol and Surface Stability of SARS-CoV-2 as Compared with SARS-CoV-1. N Engl J Med. 2020;382(16):1564–7. Epub 2020/03/18. doi: 10.1056/NEJMc2004973. PubMed PMID: 32182409; PubMed Central PMCID: PMCPMC7121658.

8. Patterson EI, Prince T, Anderson ER, Casas-Sanchez A, Smith SL, Cansado-Utrilla C, et al. Methods of Inactivation of SARS-CoV-2 for Downstream Biological Assays. J Infect Dis. 2020;222(9):1462–7. Epub 2020/08/17. doi: 10.1093/infdis/jiaa507. PubMed PMID: 32798217; PubMed Central PMCID: PMCPMC7529010.

9. Chan KH, Sridhar S, Zhang RR, Chu H, Fung AY, Chan G, et al. Factors affecting stability and infectivity of SARS-CoV-2. J Hosp Infect. 2020;106(2):226–31. Epub 2020/07/12. doi: 10.1016/j.jhin.2020.07.009. PubMed PMID: 32652214; PubMed Central PMCID: PMCPMC7343644.

10. Hellewell J, Abbott S, Gimma A, Bosse NI, Jarvis CI, Russell TW, et al. Feasibility of controlling COVID-19 outbreaks by isolation of cases and contacts. Lancet Glob Health. 2020;8(4):e488–e96. Epub 2020/03/03. doi: 10.1016/S2214-109X(20)30074-7. PubMed PMID: 32119825; PubMed Central PMCID: PMCPMC7097845.

11. Ogbunugafor CB, Miller-Dickson MD, Meszaros VA, Gomez LM, Murillo AL, Scarpino SV. The intensity of COVID-19 outbreaks is modulated by SARS-CoV-2 free-living survival and environmental transmission. medRxiv. 2020. Epub 2020/06/09. doi: 10.1101/2020.05.04.20090092. PubMed PMID: 32511513; PubMed Central PMCID: PMCPMC7273281.

12. Infection Prevention and Control of Epidemic- and Pandemic-Prone Acute Respiratory Infections in Health Care. WHO Guidelines Approved by the Guidelines Review Committee. Geneva 2014.

13. Razzini K, Castrica M, Menchetti L, Maggi L, Negroni L, Orfeo NV, et al. SARS-CoV-2 RNA detection in the air and on surfaces in the COVID-19 ward of a hospital in Milan, Italy. Sci Total Environ. 2020;742:140540. Epub 2020/07/04. doi: 10.1016/j.scitotenv.2020.140540. PubMed PMID: 32619843; PubMed Central PMCID: PMCPMC7319646.

14. Coley SE, Lavi E, Sawicki SG, Fu L, Schelle B, Karl N, et al. Recombinant mouse hepatitis virus strain A59 from cloned, full-length cDNA replicates to high titers in vitro and is fully pathogenic in vivo. J Virol. 2005;79(5):3097–106. Epub 2005/02/15. doi: 10.1128/JVI.79.5.3097-3106.2005. PubMed PMID: 15709029; PubMed Central PMCID: PMCPMC548458.

15. Richards GP. Critical review of norovirus surrogates in food safety research: rationale for considering volunteer studies. Food Environ Virol. 2012;4(1):6–13. Epub 2012/03/13. doi: 10.1007/s12560-011-9072-7. PubMed PMID: 22408689; PubMed Central PMCID: PMCPMC3284674.

16. Song H, Li J, Shi S, Yan L, Zhuang H, Li K. Thermal stability and inactivation of hepatitis C virus grown in cell culture. Virol J. 2010;7:40. Epub 2010/02/20. doi: 10.1186/1743-422X-7-40. PubMed PMID: 20167059; PubMed Central PMCID: PMCPMC2834657.

17. Sciences TAoM. Preparing for a challenging winter 2020/21. 2020.

18. Dear K, Grayson L, Nixon R. Potential methanol toxicity and the importance of using a standardised alcohol-based hand rub formulation in the era of COVID-19. Antimicrob Resist Infect Control. 2020;9(1):129. Epub 2020/08/11. doi: 10.1186/s13756-020-00788-5. PubMed PMID: 32771064; PubMed Central PMCID: PMCPMC7414286.

19. Klemes JJ, Fan YV, Jiang P. The energy and environmental footprints of COVID-19 fighting measures - PPE, disinfection, supply chains. Energy (Oxf). 2020;211:118701. Epub 2020/09/02. doi: 10.1016/j.energy.2020.118701. PubMed PMID: 32868962; PubMed Central PMCID: PMCPMC7450254.

20. Dicken RDG, T.; Perks, S. Overcoming the Regulatory Hurdles for the Production of Hand Sanitizer for Public Health Protection: The UK and US Academic Perspective. ACS Chemical Health and Safety. 2020;27:209–13. doi: 10.1021/acs.chas.0c00065.

